# The IFN-γ Paradox: Mechanistic Modeling Uncouples Cytokine Abundance from Influenza Pathogenesis

**DOI:** 10.64898/2026.11.23.768736

**Authors:** Cailan Jeynes-Smith, Amanda P. Smith, Nicole Bruce, Hai-Hsin Huang, Klaus Schughart, Meagan Herald, Amber M. Smith

**Affiliations:** Department of Pediatrics, University of Tennessee Health Science Center, Memphis, TN USA; Department of Microbiology, Immunology, and Biochemistry, University of Tennessee Health Science Center, Memphis, TN USA; Department of Mathematics, Virginia Military Institute, Lexington, VA USA; Institute of Virology Muenster, University of Muenster, Muenster, Germany

## Abstract

IFN-γ has both pathogenic and protective roles during influenza infection, yet its value as a biomarker remains unclear because both increased and decreased levels associate with greater severity. To resolve this, we developed a mechanistic mathematical model of IFN-γ and IFN-γ-producing cell kinetics. The model was calibrated using murine data and validated against immune perturbation experiments, murine transcriptomics, human influenza challenge, and critically ill patient data. Personalizing the model with human challenge data quantified interindividual heterogeneity and generating cohorts of virtual patients with varying degrees of severity predicted unmeasured dynamics and was validated with transcriptomic and proteomic data from critically ill patients. Although severe disease was associated with greater receptor-mediated IFN-γ uptake despite comparable production in non-ICU and ICU patients, early innate IFN-γ signaling was a stronger determinant. These findings explain divergent IFN-γ responses and establish a mechanistic framework for interpreting cytokine measurements.

## INTRODUCTION

Influenza viruses remain a major cause of seasonal epidemics and continue to impose a substantial global disease burden. Severe disease has been linked to robust viral replication in the lower respiratory tract and dysregulated host immune responses, which together drive acute lung injury (*1–6*). Among the cytokines implicated in influenza pathogenesis, IFN-γ plays a crucial mediator because of its antiviral activity, regulation of immune cell function, and coordination of adaptive immune responses (*1, 7–12*). However, the relationship between IFN-γ and disease severity remains difficult to interpret because cytokine concentrations reflect the combined effects of immune activation and regulatory feedback mechanisms (*7–14*). While biomarkers relating to cytokines may correlate with severity at the population level, they fail to distinguish the underlying biological mechanisms or reliably predict outcomes in individual patients.

During influenza virus infection, IFN-γ is produced by NK cells, CD4^+^ T cells, and CD8^+^ T cells (*15–17*). Through these cell populations, IFN-γ coordinates diverse aspects of the antiviral immune response by enhancing antigen presentation, promoting macrophage activation and phagocytic function, stimulating leukocyte recruitment and T cell cytotoxicity, and supporting the development of memory T cells (*18–25*). However, IFN-γ also contributes to inflammatory pathways associated with severe influenza by promoting recruitment and activation of macrophages and T cells that can exacerbate immunopathology during severe infection (*18, 26– 33*). Consequently, IFN-γ serves a dual role during influenza infection, where it is essential for promoting antiviral immunity and regulating protective immune responses, yet sustained or dysregulated IFN-γ signaling can amplify inflammation and contribute to tissue damage (*1, 18, 26–32*).

Clinical and experimental studies report conflicting associations between IFN-γ and severity, with both increased and decreased IFN-γ levels observed in severe disease across diverse patient populations and study designs (*7–12, 29*). Similarly, studies in mice have shown that reducing IFN-γ signaling through genetic deletion of IFN-γ, IFN-γ receptor deficiency, and antibody-mediated neutralization can improve disease outcomes (*1, 26, 29, 34–37*). However, these interventions do not selectively alter IFN-γ activity and can simultaneously influence viral burden, macrophage abundance, and CD8^+^ T cell cytotoxicity, among other immune processes, making it challenging to determine why IFN-γ levels are altered and whether they connect to changes in disease severity (*26, 35, 37*). This highlights a fundamental challenge in interpreting IFN-γ measurements and using cytokines to predict disease outcome because measured IFN-γ levels reflect the combined effects of cytokine production, cellular consumption, and receptor availability. A time-resolved understanding of how these processes interact is therefore essential for determining when IFN-γ promotes protection versus immunopathology.

Mechanistic computational models provide a complementary approach by integrating these interacting processes to identify the biological mechanisms underlying cytokine dynamics and their relationship to clinical outcomes. Our prior mathematical modeling studies have linked viral load and CD8^+^ T cell dynamics with immunopathology (*6*) and demonstrated that the infection remain predictable for human infection despite inter-individual heterogeneity in viral, immune, and symptom dynamics (*38*). Extending these frameworks to IFN-γ has the potential to resolve long-standing questions regarding its role in influenza pathogenesis by distinguishing the contributions of individual IFN-γ-producing cell populations, cytokine production, and receptor-mediated uptake. However, existing approaches rely on prescribed kinetics for cytokine-producing cells or are constrained only by measurements of cytokine abundance. As a result, they can reproduce observed IFN-γ trajectories without uniquely resolving the underlying relationships among immune cell dynamics, cytokine production, receptor-mediated uptake, and disease severity (*19*).

Here, we developed a mechanistic mathematical model that integrates time-resolved measurements IFN-γ and IFN-γ^+^ NK cells, CD4^+^ T cells, and CD8^+^ T cells, allowing cytokine production to be dynamically distinguished from receptor-mediated uptake while quantifying the contributions of individual immune cell populations to observable IFN-γ dynamics. Model-derived estimates of IFN-γ production and receptor-mediated uptake were independently supported by transcriptomic data, establishing important quantitative links between experimentally measurable gene expression and underlying cytokine dynamics. The results were validated in immune perturbation experiments and demonstrate that similar IFN-γ concentrations can arise from fundamentally different immune states, providing a mechanistic explanation for the conflicting associations between IFN-γ and influenza disease severity reported across studies. Applying the model to human influenza further quantified interindividual heterogeneity in IFN-γ responses and revealed that severe disease can be associated with greater receptor-mediated uptake of IFN-γ despite comparable levels of IFN-γ production between non-ICU and ICU patients. Although neither IFN-γ abundance nor inferred receptor-mediated uptake reliably distinguished severity on their own, early IFN-γ^+^ NK-cell responses emerged as the strongest discriminator of severity, highlighting a previously underappreciated role for early innate IFN-γ signaling in shaping influenza outcomes.

## RESULTS

### Temporal Divergence Between IFN-γ Levels, IFN-γ^+^ Cell Counts, and Fluorescence Intensity

Following infection of mice with 75 TCID_50_ influenza A/Puerto Rico/8/34 (PR8), IFN-γ remained at baseline until 2 days post-infection (dpi) before exhibiting biphasic dynamics (Fig. 1A). The early IFN-γ peak at 3 dpi coincided with IFN-γ^+^ NK cells, whereas the later peak at 6 dpi coincided with IFN-γ^+^ CD4^+^ and CD8^+^ T cells. Consistent with previous reports (*39*), no other immune or epithelial cell populations were detected as sources of IFN-γ. Notably, the IFN-γ^+^ NK cell dynamics differed markedly from those of the broader NK cell population, which peaked at 7 dpi (Fig. S1), indicating that the total NK cell abundance is a poor surrogate for IFN-γ production. In contrast, IFN-γ^+^ CD4^+^ and CD8^+^ T cells exhibited similar temporal dynamics as each other and their total population, with expansion beginning at 4-5 dpi and IFN-γ^+^ CD4^+^ T cells expanding more rapidly than IFN-γ^+^ CD8^+^ T cells (5 dpi, *P* = 0.02; Fig. 1A). The decline in IFN-γ^+^ NK cells at 4-5 dpi paralleled the expansion of IFN-γ^+^ T cells, suggesting a coordinated transition from innate to adaptive sources of IFN-γ production.

**Fig. 1.**
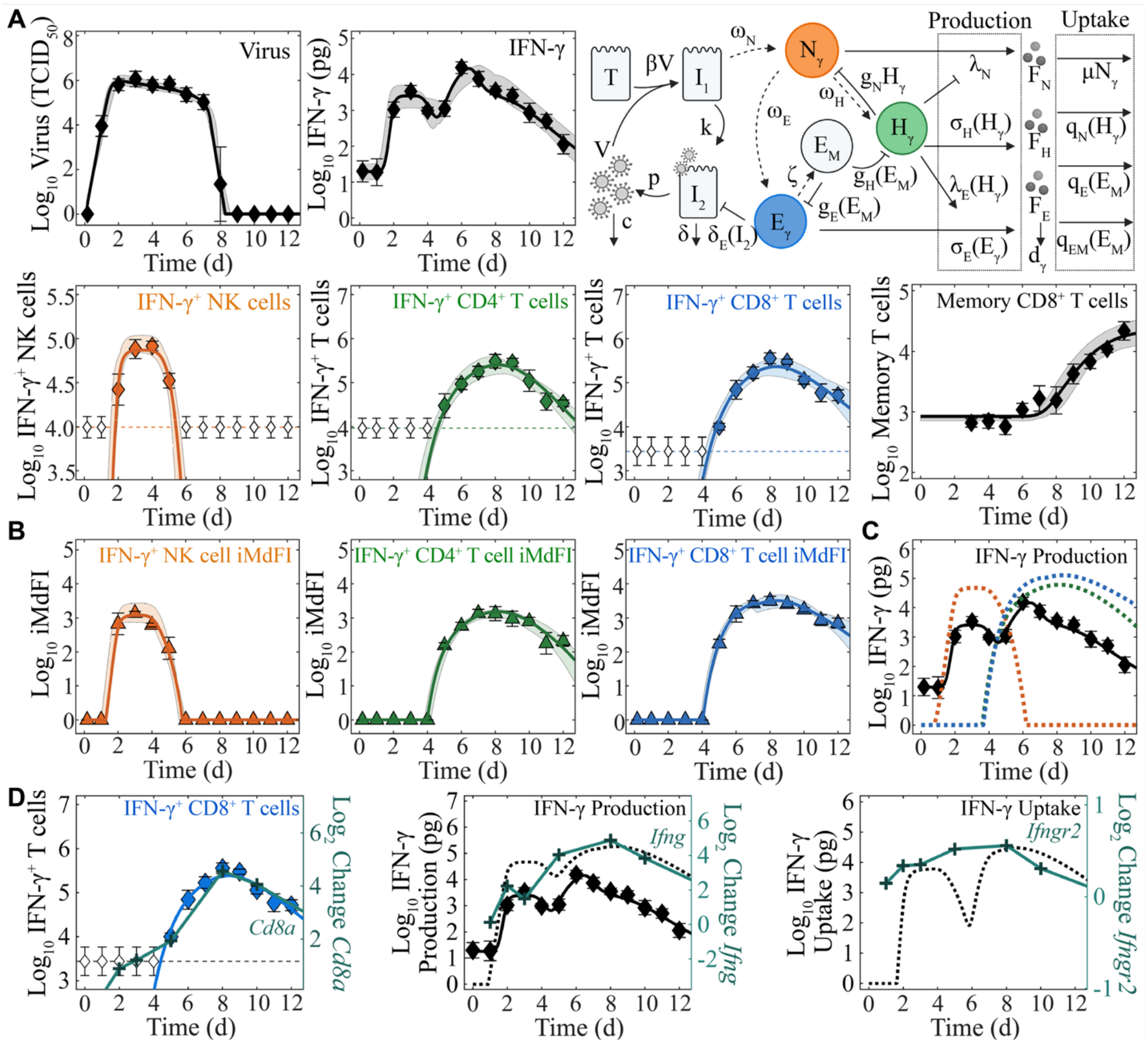
Dynamics of IFN-γ-producing immune cells and mechanistic modeling of IFN-γ regulation during influenza. (**A**-**B**) Fit of the mathematical model to experimental data, including (**A**) viral loads, IFN-γ, CD43^-^CD127^+^CD103^+^CD69^+^CD8^+^ memory T cells, IFN-γ^+^ NK cells, IFN-γ^+^ CD4^+^ T cells, IFN-γ^+^ CD8^+^ T cells, and (**B**) their integrated median fluorescence intensities, from mice infected with influenza A/PR8. The underlying model structure, including IFN-γ-producing cell dynamics and regulation of IFN-γ production and receptor-mediated uptake, is depicted in the schematic and Equations (1)-(15). Model parameter descriptions, and best fit estimates and 95% confidence intervals are listed in Table S1. (**C**) Model-inferred contributions of IFN-γ production by IFN-γ^+^ NK cells (dashed orange line), IFN-γ^+^ CD4^+^ T cells (dashed green line), and IFN-γ^+^ CD8^+^ T cells (dashed blue line), illustrating the divergence between total production and observed IFN-γ levels attributable to receptor-mediated IFN-γ uptake. (**D**) Consistency of published transcriptomic data from C57BL/6 mice **(*40*)** for *Cd8a* (teal plus signs) with IFN-γ^+^ CD8^+^ T cells (blue diamonds), *Ifng* (teal plus signs) with IFN-γ production (black dashed line), and *Ifngr2* (teal plus signs) with IFN-γ receptor-mediated uptake (black dashed line). The delay between *Ifngr2* and inferred IFN-γ uptake suggests receptor availability is a limiting factor.

The amount of IFN-γ produced per cell varied dynamically throughout the infection as indicated by the integrated median fluorescence intensity (iMdFI) of IFN-γ from producing cells (Fig. 1B). IFN-γ^+^ NK cells reached fluorescence intensities comparable to IFN-γ^+^ T cells, and IFN-γ^+^ CD8^+^ T cells consistently exhibited higher fluorescence intensities than CD4^+^ T cells (6-12 dpi; all *P* < 0.01). However, fluorescence intensities were uncoupled from both cell count and IFN-γ levels. For example, IFN-γ^+^ NK cell fluorescence intensity remained relatively constant between 2 and 4 dpi despite a 0.5 log_10_ increase in IFN-γ^+^ NK cell numbers from 2 to 3 dpi. Likewise, peak IFN-γ fluorescence intensity in T cells (7-8 dpi) did not coincide with the second IFN-γ peak (6 dpi). These observations indicate that measured IFN-γ concentrations cannot be inferred directly from abundance of IFN-γ producing cells and instead reflect the combined effects of dynamic changes in cell number, cell-specific functional responsiveness, and downstream cytokine handling.

### Opposing Production and Receptor-Mediated Uptake Shape IFN-γ Dynamics

The temporal discordance between IFN-γ, IFN-γ^+^ cell counts, and their intracellular IFN-γ fluorescence suggested that observable cytokine dynamics could not be explained by changes in immune populations alone. To resolve these relationships, we developed a mechanistic model of IFN-γ regulation that explicitly describes the regulatory dynamics of IFN-γ-producing cells together with cytokine production and receptor-mediated uptake (Equations (1)-(15); Fig. 1A, 1B). The model accurately recapitulated the observed IFN-γ, IFN-γ^+^ NK cell, IFN-γ^+^ CD4^+^ T cell, and IFN-γ^+^ CD8^+^ T cell dynamics, including intracellular fluorescence intensity, enabling the relative contributions of individual immune cell populations to IFN-γ production and receptor-mediated uptake to be quantified (Fig. 1C). The model further revealed that detectable IFN-γ is shaped by opposing effects of cytokine production and receptor-mediated uptake together with cell type-specific regulatory interactions among NK cells and T cells, providing a mechanistic explanation for the experimentally observed temporal discordance.

Analysis of the inferred regulatory interactions showed that both autocrine and paracrine signaling contributed to the observed cytokine dynamics. NK cell-derived IFN-γ was predominantly taken up by CD4^+^ T cells, whereas T cell-derived IFN-γ was internalized by effector and memory T cells. The model further predicted reciprocal regulation between IFN-γ^+^ T cells and IFN-γ^+^ NK cells, with CD4^+^ T cells suppressing IFN-γ activity in NK cells while enhancing IFN-γ production by IFN-γ^+^ CD8^+^ T cells.

### Mechanistic Modeling Links Transcriptomics to IFN-γ Production and Signaling

Transcriptomic data are increasingly available, yet translating changes in gene expression into cytokine production and signaling remains a major challenge. To determine whether transcriptomic measurements could inform otherwise unobservable IFN-γ dynamics, we compared the respective published transcriptomic data from influenza-infected C57BL/6 mice (*40*) with model-derived estimates of IFN-γ-producing cell populations, cytokine production, and receptor-mediated uptake. *Cd4a, Cd8a*, and *Cd8b1* expression closely tracked the dynamics of IFN-γ^+^ CD4^+^ and CD8^+^ T cells, respectively (Fig. 1D; Fig. S2), supporting their use as proxies for activated T cell populations within the modeling framework. In contrast, NK cell-associated transcripts (*Klra1*-*Klra23, Klrb1a, Klrb1c, Klrb1f, Klre1, Klri2, Ncr1*, or *Tmsb15a*) did not reflect IFN-γ^+^ NK cells, consistent with the phenotypic divergence between the total and IFN-γ producing NK cell populations (Fig. S1; Fig. S2).

Importantly, *Ifng* expression closely matched the model-predicted IFN-γ production but not the observed IFN-γ protein levels (Fig. 1D). Conversely, *Ifngr2* expression closely followed the inferred receptor-mediated uptake of IFN-γ, albeit with a ∼1 day delay (Fig. 1D). This alignment indicates that expression of the receptor is the primary limit on bound IFN-γ for most of the infection, whereas IFN-γ availability likely becomes limiting between 4 and 7 dpi. These findings establish a mechanistic link between transcriptomic measurements and the underlying processes governing cytokine dynamics, demonstrating that transcriptomics can inform cytokine production and receptor-mediated uptake rather than cytokine abundance alone.

### CD8^+^ T cell Depletion Reduces Total Bound IFN-γ, Increasing Free Levels

Because CD8^+^ T cells become the dominant producers of IFN-γ during infection resolution (7-9 dpi), we examined how their depletion alters IFN-γ production and receptor-mediated uptake. CD8^+^ T cell depletion resulted in transiently lower IFN-γ early after infection (2 dpi [*P* = 0.05]; 4 dpi [*P* = 0.01]) but higher IFN-γ (8 dpi [*P* < 10^-3^]) later in infection, despite reduced viral loads and improved disease severity (*6*) (Fig. 2A, 2C). The depletion also induced early innate immune depletion and activation, characterized by rapid recovery of NK cells (0 dpi, *P* = 0.04; 2 dpi, *P* = 0.1; Fig. S3A; Fig. 2A) together with reduced CD4^+^ T cells early in infection (2 dpi, *P* = 0.04) and increased CD4^+^ T cells late in infection (6 and 10 dpi, *P* = 0.06; Fig. S3B; Fig. 2A).

**Fig. 2.**
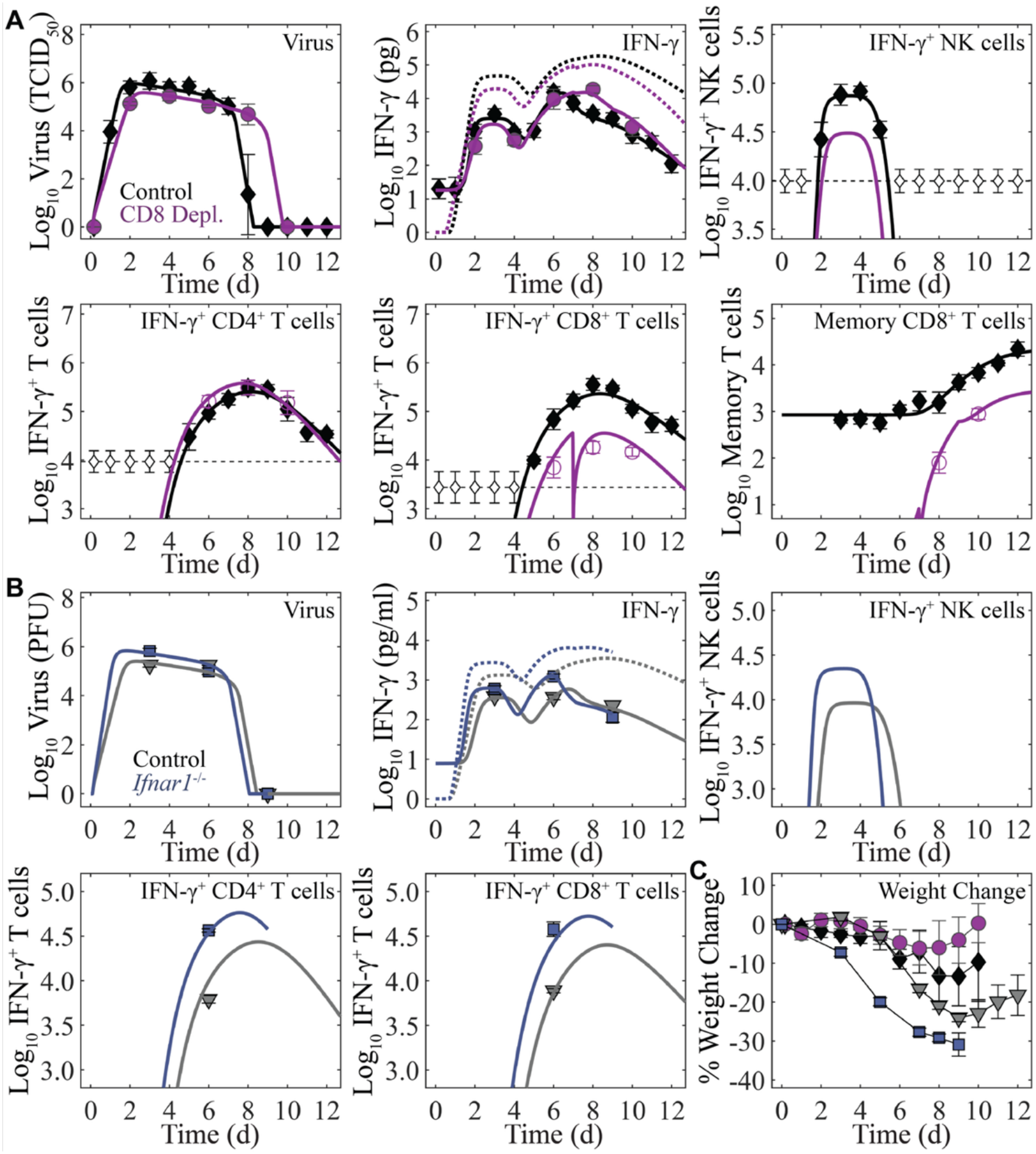
Mechanistic Model Explains the Emergence of Elevated IFN-γ Despite Differential Severity in CD8^+^ T cell-Depleted and *Ifnar1*^-/-^ Mice. (**A**) Model fits to viral loads (solid markers), IFN-γ (solid markers), and predicted changes to IFN-γ^+^ CD4^+^ and CD8^+^ T cells and to CD43^-^ CD127^+^CD103^+^CD69^+^CD8^+^ memory T cells (open markers) from BALB/c control (black) and CD8^+^ T cell-depleted mice (magenta) infected with influenza A/PR8 (*6*). CD8^+^ T cell depletion resulted in a lower infection size (*T*_0_), altered IFN-γ^+^ cell activation timing (τ_*N*_, τ_*H*_, and τ_*E*_), and an increased CD4^+^ T cell activation rate (ω_*H*_). Inferred IFN-γ production for the depletion group remained below that of the control group despite the elevated CD4^+^ T cell activation rate (dotted line). (**B**) Increased viral loads, IFN-γ, CD4^+^ T cells, and CD8^+^ T cells from C57BL/6 control (grey) and *Ifnar1*^-/-^ (blue) mice infected with influenza A/FM (*36*) were explained by a higher infection size (*T*_0_) in *Ifnar1*^-/-^ mice. (**C**) Weight loss in *Ifnar1*^-/-^ mice was higher than in control C57BL/6 mice and lower in CD8^+^ T cell-depleted mice than in BALB/c mice.

To determine how these changes gave rise to the observed IFN-γ dynamics, the model was refit to the depletion data (Fig. 2A). The model predicted earlier activation of IFN-γ^+^ NK cells (by ∼10 h) in addition to earlier (by 9 h) and faster (3.8x) activation of IFN-γ^+^ CD4^+^ T cells (Table S2), confirming early immune activation, whereas the delayed arrival of IFN-γ^+^ CD8^+^ T cells (by 19 h) and altered downregulation of IFN-γ^+^ T cells (∼10x lower half-saturation constants) reflected the depletion schedule. Importantly, no changes to IFN-γ production, receptor-mediated uptake, cytotoxicity, or other functional parameters were required. Instead, the altered IFN-γ dynamics were explained by the reduced number of infected cells and loss of CD8^+^ T cells. Fewer infected cells in the CD8^+^ T cell depleted groups naturally led to lower predicted numbers of IFN-γ^+^ NK cells and subsequently less NK cell-derived IFN-γ production. Although CD4^+^ T cells were more abundant, it did not compensate for the reduced contribution of IFN-γ from CD8^+^ T cells (Fig. 2A, dotted lines). Rather, the elevated IFN-γ levels observed late in infection (8 dpi) resulted from substantially reduced receptor-mediated uptake due to the depletion of CD8^+^ T cells. These findings demonstrate that elevated IFN-γ does not necessarily indicate increased cytokine production. Instead, reduced receptor-mediated uptake can increase measurable IFN-γ despite lower overall cytokine production, emphasizing the importance of accounting for both processes when interpreting cytokine measurements.

### Increased Viral Burden Drives Elevated IFN-γ in *Ifnar1*^-/-^ Mice

To determine whether our model could distinguish changes in IFN-γ dynamics arising from increased viral burden rather than altered cytokine regulation, we analyzed data from mice lacking IFN-α/β receptor signaling (*36*). In contrast to CD8^+^ T cell depletion, which reduced viral burden, loss of type I IFN signaling impairs antiviral control, resulting in higher viral loads, increased numbers of IFN-γ^+^ CD4^+^ and CD8^+^ T cells, elevated IFN-γ, and greater disease severity (Fig. 2B, 2C) (*36*). The model accurately recapitulated the infection dynamics in C57BL/6 control mice by reducing the rate of NK cell-derived IFN-γ uptake (by 0.1x; Fig. 2B; Table S3), a difference consistent in direction with the elevated IFN-γ responses reported in BALB/c mice (*35, 41*). After accounting for mouse strain-specific differences, the only additional change required to reproduce the *Ifnar1*^-/-^ data was a 2.9x increase in infection size (Fig. 2B; Table S3), reflecting the loss of IFN-α/β-mediated antiviral control. The effects of this larger infected cell population propagated through the immune response, driving the observed increases in IFN-γ and IFN-γ^+^ T cells and a predicted 3x increase in IFN-γ^+^ NK cells. Although total IFN-γ production increased (Fig. 2B, dotted lines), receptor-mediated uptake also increased because of the expansion of IFN-γ-responsive immune cells, resulting in similar measured IFN-γ at 3 and 9 dpi. Importantly, no changes to intrinsic IFN-γ production or receptor-mediated uptake rates, or other host-pathogen interaction parameters, were required to reproduce these dynamics, indicating that the elevated IFN-γ response in *Ifnar1*^-/-^ mice emerged from increased viral burden and its downstream immune consequences rather than altered IFN-γ kinetics.

### Individualized IFN-γ Dynamics During Human Influenza Challenge

To determine whether the model generalized to human influenza infection, we analyzed data from a human influenza challenge study (*42*). In our previous work, we showed that interindividual differences in viral loads and CD8^+^ T cell dynamics in serum were explained by individualized rates of infection and CD8^+^ T cell expansion after accounting for differences in infection size (*38*). Extending this framework to include IFN-γ accurately recapitulated the observed IFN-γ, viral loads, and CD8^+^ T cell trajectories across individuals (Fig. 3A; Fig. S4; Table S4). The model additionally predicted heterogeneous IFN-γ^+^ NK cells with peaks spanning ∼1.5 log_10_ cells (Fig. 3A), despite these cells not being measured.

**Fig. 3.**
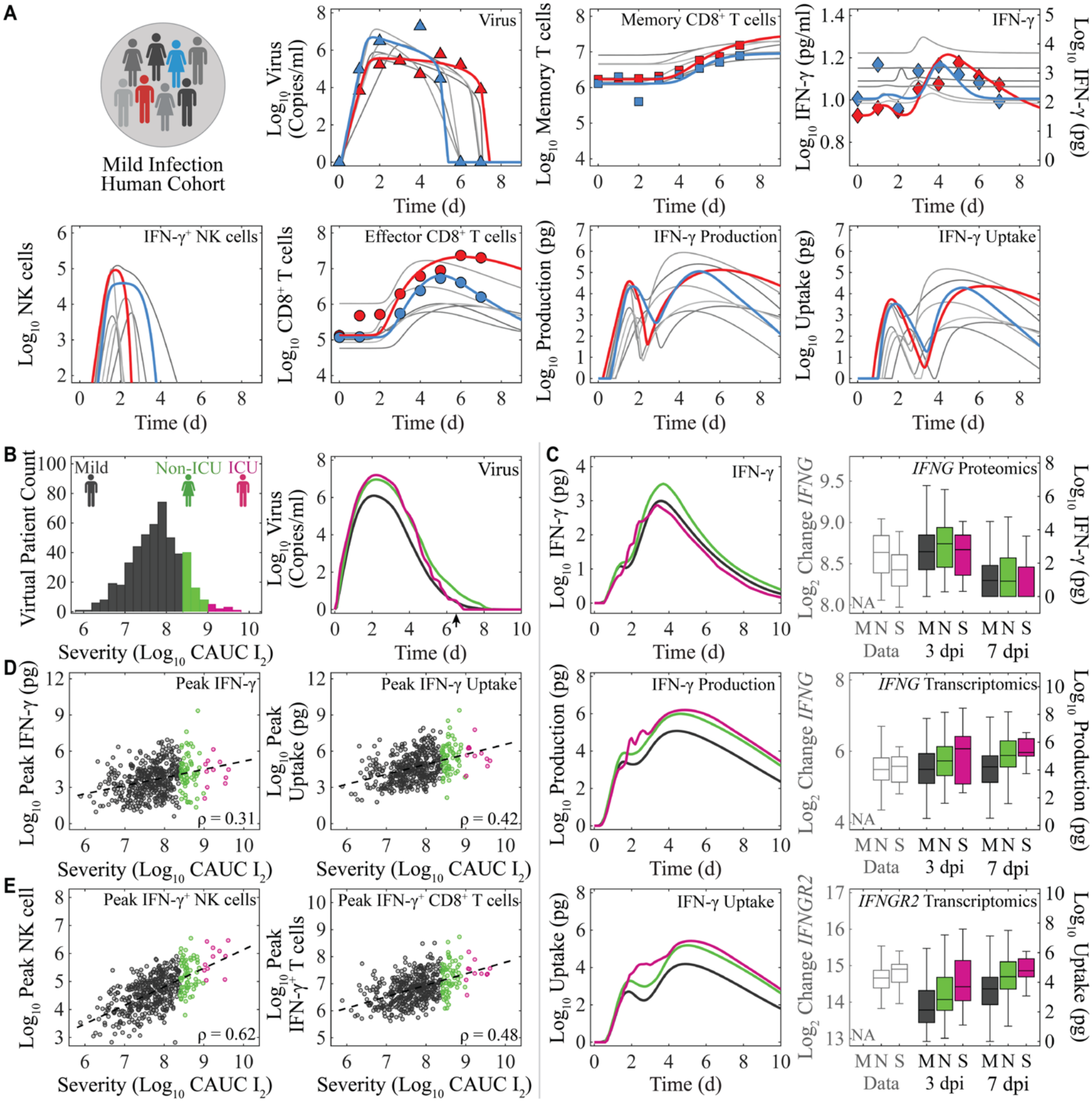
Virtual Patient Simulations Identify Receptor-Mediated Uptake and IFN-γ^+^ NK cells as Biomarkers of Influenza Disease Severity. (**A**) Model simulations of viral load, effector (CD38^+^CD45RA^+^CD27^-^CCR7^-^) and memory (CD38^+^CD45RA^-^CD27^+^) CD8^+^ T cells, and IFN-γ data from a human challenge study (*38, 42*). Data from two representative patients are shown (red and blue) together with model trajectories for the remaining participants (gray; all fits are shown in Fig. S4) to illustrate inter-individual heterogeneity. (**B**) Characterization of mild (black; *M*), moderate (green; non-ICU; *N*), and severe (magenta; ICU; *S*) virtual patient cohorts with representative viral load trajectories. (**C**) Simulated IFN-γ concentrations, IFN-γ production, and receptor-mediated IFN-γ uptake across severity cohorts, with proteomic (IFNG) and transcriptomic (*IFNG* and *IFNGR2*) measurements from influenza-infected non-ICU and ICU patients (*9, 10*). (**D**-**E**) Peak simulated IFN-γ concentrations, receptor-mediated IFN-γ uptake (**D**), IFN-γ^+^ NK cells, and IFN-γ^+^ CD8^+^ T cells (**E**) and the associated Pearson’s linear correlation coefficient, ρ, with severity. IFN-γ^+^ NK cells distinguished all three severity groups, whereas peak IFN-γ concentrations, IFN-γ uptake, and IFN-γ^+^ CD8^+^ T cell abundance differentiated severity less effectively with significant overlap across all cohorts.

Across individuals, inferred IFN-γ receptor-mediated uptake rates increased in proportion to T cell activation rates, consistent with the role of NK cell-derived IFN-γ in promoting immune cell activation (*16, 20, 25*) (Fig. 3A; Fig. S4; Table S4). Similarly, the rates of IFN-γ uptake by CD8^+^ T cells scaled with their efficacy, supporting the role of IFN-γ in CD8^+^ T cell effector function (*1, 21, 22, 25*). Finally, the rates of IFN-γ production by CD4^+^ and CD8^+^ T cells scaled with the rate of memory CD8^+^ T cell formation, consistent with the established role of IFN-γ in memory T cell development (*20–24*). Despite these relationships, IFN-γ production and receptor-mediated uptake exhibited disproportionately greater inter-individual variability than immune cell abundances (Fig. 3A), suggesting that heterogeneity in IFN-γ dynamics is driven primarily by differences in its regulation rather than cell number.

### Higher IFN-γ^+^ NK cells and IFN-γ Uptake Corresponds with Influenza Severity in Humans

To further investigate how heterogeneity in IFN-γ responses relates to influenza disease severity, we generated 500 virtual patients and classified them as having mild (*M*), moderate (non-ICU; *N*), or severe (ICU; *S*) influenza based on their infection size (Fig. 3B). The immune kinetics for each severity cohort over the span of the infection can be found in Fig. S5. To evaluate these predictions, we compared simulated IFN-γ concentrations, IFN-γ production, and receptor-mediated IFN-γ uptake across severity groups with independent proteomic (IFNG) and transcriptomic (*IFNG* and *IFNGR2*) data from influenza-infected non-ICU and ICU patients (*9, 10*). The virtual cohort closely recapitulated the trends observed in the clinical data. Both the human cohort and the virtual patients predicted to have severe disease exhibited modestly lower IFN-γ protein levels than the corresponding non-ICU group (Fig. 3C). Estimated IFN-γ production increased with disease severity in the virtual cohort but did not differ significantly between the groups, consistent with the *IFNG* transcriptomic data. In contrast, IFN-γ uptake increased in the severe virtual cohort and was similarly associated with higher *IFNGR2* expression in ICU patients. Notably, these results indicated that hospitalized patients maintained higher IFN-γ production and uptake despite having circulating IFN-γ concentrations that were comparable to, or lower than, those of patients with milder disease. This disconnect suggests that circulating IFN-γ alone does not reflect the underlying cytokine dynamics, which are determined by the balance between IFN-γ production and receptor-mediated uptake. Consistent with this finding, peak circulating IFN-γ was the least informative discriminator of disease severity in the virtual cohort (Fig. 3D), whereas IFN-γ^+^ NK cells uniquely distinguished mild, non-ICU, and ICU cohorts (Fig. 3E). Together, these findings identify IFN-γ^+^ NK cells and the associated differences in IFN-γ uptake as more informative indicators of disease severity than circulating IFN-γ concentrations alone.

## DISCUSSION

Our study provides a mechanistic explanation for the conflicting associations between IFN-γ and influenza disease severity reported across experimental and clinical studies (*1, 7–12*). By integrating mechanistic modeling with murine experiments, human influenza challenge data, and independent clinical transcriptomic and proteomic datasets, we demonstrated that IFN-γ concentrations are predictable and determined by the balance between cytokine production and receptor-mediated uptake rather than production alone. Consequently, similar IFN-γ concentrations can arise from fundamentally different underlying immune dynamics, limiting the utility of IFN-γ or other cytokines as standalone biomarkers of disease severity without the use of mechanistic models. Rather than viewing IFN-γ as simply protective or pathogenic, our findings suggest that its relationship with disease severity depends on the dynamic interplay between cytokine production, cellular uptake, and the responding immune populations.

A major methodological advance of this study is the integration of cell type-specific IFN-γ production into a mechanistic modeling framework. By combining intracellular cytokine measurements with mechanistic modeling, we estimated IFN-γ production from individual immune cell populations and established a quantitative relationship between these estimates and transcriptomic measurements. Although intracellular cytokine measurements provide valuable experimental insight by enabling resolution of cell-specific IFN-γ production, they are rarely available in human studies. The quantitative correspondence between model-derived IFN-γ production and transcript abundance therefore provides a practical framework for interpreting increasingly common transcriptomic datasets in terms of the underlying cytokine biology, extending mechanistic insight from experimental systems to human disease.

Beyond estimating cytokine production, our results demonstrate that cytokine concentrations should not be interpreted as direct measures of activity. Instead, measured IFN-γ represents the net outcome of cytokine production and receptor-mediated uptake, both of which change dynamically throughout infection. This distinction provides a mechanistic explanation for why studies have reported both increased and decreased IFN-γ concentrations in severe influenza (*7–13*) despite IFN-γ’s established roles in antiviral immunity and immunopathology (*18–32*). More broadly, these findings suggest that cytokine abundance alone may be an incomplete measure of immune function and highlight the value of mechanistic models for disentangling the competing biological processes that shape measurable cytokine responses.

Our findings also identify early innate IFN-γ signaling as a potentially underappreciated determinant of influenza disease severity. Across multiple experimental perturbations and in the human patient simulations, early IFN-γ^+^ NK cell responses consistently tracked with subsequent disease severity, whereas later circulating IFN-γ concentrations were comparatively uninformative. These observations suggest that the timing and cellular source of IFN-γ may be more important than its abundance alone. Early IFN-γ signaling has been linked to activation of inflammatory macrophages and the subsequent development of immunopathology (*9, 10*), providing a plausible biological mechanism through which relatively small differences in early innate responses could shape later disease outcomes. While these downstream pathways require further experimental validation, our results support a model in which early innate IFN-γ signaling establishes the inflammatory trajectory that ultimately determines disease severity.

These findings have broader implications for the development and interpretation of immune biomarkers, particularly as IFN-γ production and receptor-mediated uptake varies throughout infection and between individuals. Most clinical studies rely on measurements of circulating cytokines obtained at a single time point, implicitly assuming that cytokine abundance reflects cytokine activity. Our results challenge this assumption by demonstrating that comparable circulating IFN-γ concentrations can arise from markedly different underlying production and uptake dynamics. Accordingly, biomarkers that incorporate the cellular processes regulating cytokine signaling, or mechanistic models capable of inferring these latent processes, may provide more informative predictors of disease severity than cytokine concentrations alone. More broadly, this framework offers a general strategy for interpreting cytokine measurements across infectious and inflammatory diseases where production and receptor-mediated uptake occur simultaneously.

Predictive models that apply across multiple infection scenarios, like the one presented here, have important implications for immune digital twins and other patient-specific precision medicine approaches. A central challenge for digital twin technologies is that many of the biological processes driving disease progression cannot be measured directly in individual patients. In addition, mechanistic models are often custom built for a single scenario, which limits their broader applicability (*43*). Fitting our model to human data and simulating virtual patients allowed us to explore how variability in host responses shapes infection outcomes. Our results also show that IFN-γ concentrations alone provide an incomplete picture and that integrating transcriptomic and proteomic signatures can help bridge this gap. However, two key challenges remain. First, identifying patient-specific parameters is difficult when data are sparse, and distinguishing which parameters must be individualized from those that can be fixed at population values will be critical for reliable predictions. Second, it is not yet clear which model-derived metrics best predict severity. Quantities, such as infected cell burden (*6, 44*) or cumulative inflammation (*45*), may ultimately prove more informative than individually measured biomarkers for stratifying patients, predicting disease trajectories, and identifying optimal therapeutic interventions (*43*).

This study has limitations that also suggest directions for future work. Our model focuses on the dominant IFN-γ-producing and responding immune cell populations supported by the available data and does not explicitly represent additional IFN-γ-responsive cells, including macrophages, fibroblasts, and epithelial cells, which have established roles in influenza immunopathology and tissue repair (*1, 19, 20, 22–24, 27, 28, 46–48*). This may be reflected in the nonlinearity of the receptor-mediated uptake terms or changing IFNGR2 expression. Incorporating additional populations will enable investigation of how cell type-specific IFN-γ signaling contributes to inflammation, viral clearance, and tissue repair throughout infection, particularly with respect to IFN-γ-dependent activation of inflammatory CCR2 macrophages (*26*). Likewise, extending the framework to include additional cytokines and their interactions will provide a more comprehensive representation of the immune regulatory networks governing influenza pathogenesis. This will be critical to developing computational biomarkers that can better stratify simulated patients by severity, where pulmonary inflammation is correlated to macrophages and neutrophils (*6, 44, 45*). A challenge faced in this study was extending our calibrated model results to human peripheral blood and serum, where cytokines exist in low concentrations and my not be representative of the pulmonary infection. Although the inferred cytokine production and uptake dynamics were supported by independent transcriptomic and proteomic datasets, future studies integrating mechanistic modeling with emerging single-cell and spatial profiling technologies could enable more direct validation and refinement of these inferred immune processes.

Our study demonstrates that interpreting cytokine biology requires moving beyond measurements of cytokine abundance alone. By integrating mechanistic modeling with experimental and clinical data, we showed that circulating IFN-γ reflects the dynamic balance between cytokine production and receptor-mediated uptake, providing a mechanistic explanation for the conflicting relationships between IFN-γ and influenza disease severity reported across studies. This framework enables otherwise inaccessible aspects of cytokine biology, including cell-specific production and cytokine uptake, to be inferred from experimentally measurable data while identifying early innate IFN-γ signaling as a potential determinant of disease severity. More broadly, these findings establish a generalizable strategy for linking multi-omic measurements with mechanistic models to improve the interpretation of immune responses and accelerate the development of predictive biomarkers and therapeutic strategies for influenza and other inflammatory diseases.

## MATERIALS AND METHODS

### Mice

Female BALB/c mice (6-week-old) were obtained from Charles River Laboratories (Wilmington, Massachusetts) and housed in groups of five in 38.2cm x 19.4cm x 13.0cm solid-bottom polysulfone individually ventilated cages. Rooms used for housing mice were maintained on a 12:12-hour light:dark cycle at 22±2°C with 50% humidity in a biosafety level 2 facility. Before inclusion in the experiments, mice were allowed at least 7 days to acclimate to the animal facility, such that they were 7 weeks old at the time of infection. Teklad LM-485 Mouse/Rat Sterilizable Diet (Envigo, Indianapolis, IN) and autoclaved water were available ad libitum. All experiments were performed under approved protocols and in accordance with the guidelines set forth by the Animal Care and Use Committee at UTHSC.

### Infection Experiments

Mice were intranasally inoculated with 75 TCID_50_ influenza A/Puerto Rico/8/34 (H1N1) diluted in 100 μl of sterile PBS. The viral infectious dose (TCID_50_) was determined by interpolation using the Reed and Muench method with serial dilutions of virus on Madin-Darby canine kidney (MDCK) cells. In a subset of animals, CD8^+^ T cells were depleted by IP injection at days -2, 0, 3, and 7 pi with 100 mg of the rat anti-CD8a antibody (clone 2.43) that was purified from ATCC hybridoma (per manufacturer instructions) in 250 ml of PBS as previously described(*6*). Mice were weighed at the onset of infection and each subsequent day to monitor illness and mortality, and euthanized if they became moribund or lost 30% of their starting body weight. All experiments were repeated once to ensure reproducibility, with two complete experiments (8-10 animals per time point) used for these studies. Lungs were collected and processed for immune cell isolation as previously described (*6, 44*).

### Viral Titers

Viral titers were obtained using serial dilutions on MDCK monolayers and normalized to the total volume of lung homogenate supernatant as previously described (*44*).

### Flow Cytometric Analyses

Flow cytometry (ZE5 Cell Analyzer, Bio-Rad, Hercules, CA) was performed on single cell suspensions after 20 min incubation in 200 μl of Fc block (1:2 human-γ globulin or 1:200 TruStain FcX, Biolegend) and 1:1000 viability dye (Zombie Fixable Viability Kit, Biolegend), followed by 30 min surface staining with anti-mouse antibodies and fixation and permeabilization for 20 min (BD Cytofix/Cytoperm, BD Biosciences), followed by intracellular staining for 30 min, where indicated.

For total cell quantification, single cell suspensions were stained for flow cytometric analysis as previously described using antibodies: CD11c (eFluor450, clone N418, eBioscience), CD11b (Alexa700, clone M1/70, BD Biosciences), Ly6G (PerCp-Cy5.5, clone 1A8, Biolegend), Ly6C (APC, clone HK1.4, eBioscience), F4/80 (PE, clone BM8, eBioscience), CD3e (PE-Cy7, clone 145-2C11, BD Biosciences or BV785 Biolegend), CD4 (PE-Cy5, clone RM4-5, BD Biosciences), CD8a (BV605, clone 53-6.7, BD Biosciences), CD49b (APC-Cy7, Biolegend or APC-e780, clone DX5, Affymetrix Inc), and MHC-II (FITC, clone M5/114.15.2, eBioscience). Viable cells were gated from a forward scatter/side scatter plot and singlet inclusion. Following neutrophil exclusion (Ly6G^hi^), NK cells were gated as CD3e^-^CD49b^+^, CD4^+^ T cells as CD3e^+^CD49b^-^CD4^+^CD8^-^, and CD8^+^ T cells as CD3e^+^CD49b^-^CD4^-^CD8^+^.

To quantify IFN-γ^+^ T cells, antibodies CD3e (BV786, clone 145-2C11, Biolegend), CD4 (FITC, clone RM4-5, Biolegend), CD8α (BV605, clone 53-6.7, BD Biosciences), B220 (APCe780, clone RA3-6B2, eBioscience), CD49b (APCe780, clone DX5, eBioscience), CD62L (PE-Cy7, clone MEL-14, Biolegend), CD69 (PE-Dazzle594, clone H1.2F3, Biolegend), CD25 (Alexa700, clone PC61, Biolegend), CD43 (APC, clone 1B11, Biolegend), CD127 (PE-Cy5, clone A7R34, Biolegend), and CD103 (BV711, clone 2E7, Biolegend) were used, followed by intracellular staining with IFN-γ (PE, clone XMG1.2, eBioscience). CD45 (PerCP, clone 30-F11, Biolegend) was included as either an *in vivo* label during retro-orbital injection or in cell surface staining. Viable CD4^+^ T cells were identified as CD3e^+^CD49b^-^B220^-^CD4^+^CD8^-^ and CD8^+^ T cells as CD3e^+^CD49b^-^B220^-^CD4^-^CD8^+^. In a separate panel, NK cells were quantified using antibodies CD45 (PerCP, clone 30-F11, Biolegend), CD3e (BV786, clone 145-2C11, Biolegend), CD4 (FITC, clone RM4-5, Biolegend), CD8α (BV605, clone 53-6.7, BD Biosciences), CD49b (APCe780, clone DX5, eBioscience), CD314 (NKG2D; BV711, clone CX5, BD Bioscience), CD69 (APC, clone H1.2F3, Biolegend), and CD11b (V500, clone M1/70, BD Biosciences), followed by IFN-γ staining as above. Viable NK cells were identified as CD45^+^CD3e^-^CD49b^+^, with antibodies against CD8, CD314, CD69, and CD11b used to confirm classification and assess activation. In addition, in collagenase digested lungs, the total CD45^-^ fraction in the NK cell panel was assessed for expression of intracellular IFN-γ.

IFN-γ^+^ gates were placed based on down-sampled populations (3000 events) in naïve-matched controls. The integrated median fluorescence intensity (iMdFI) was calculated as ([MdFI IFNγ^+^ - MdFI internal IFN-γ^-^] x % IFN-γ^+^ sample) - ([MdFI IFN-γ^+^ - MdFI internal IFN-γ^-^] x % IFN-γ^+^ naive). Samples were identified as below the limit of detection (LOD) when both the number of IFN-γ^+^ events in repeatedly sampled gates (2000 repeats) and the change in MdFI were less than or equal to the matched naïve control that was processed, stained, and acquired on the same day as infected animals.

Because IFN-γ^+^ cells were not quantified in the CD8^+^ T cell depletion experiment (*6*), the numbers of IFN-γ^+^ CD4^+^ and CD8^+^ T cells were inferred by applying experimentally determined proportions of IFN-γ^+^ cells within the total CD4^+^ and CD8^+^ T cell populations to the corresponding T cells counts in the depletion experiment. In contrast, IFN-γ^+^ NK cells were not imputed because the IFN-γ^+^ NK cell population was not representative of the total NK cell population, preventing reliable estimation from NK cell counts.

For all experiments, the absolute numbers of cells were calculated based on viable events analyzed by flow cytometry as related to the total number of viable cells per sample. Any value below the LOD was set to the mean value of the naïve controls. Because intracellular staining alters the viability of some cell populations, we used the kinetics of the total numbers of cells from experiments as in our prior work. All data with intracellular IFN-γ staining were analyzed using FlowJo 10.8.1 (Tree Star, Ashland, OR).

### Transcriptomics Data from Influenza Infection in Mice

We used published transcriptomic data (*Ifng, Ifngr2, Cd8a, Cd8b1, Cd4a, Klra1*-*Klra23, Klrb1a, Klrb1c, Klre1, Klrb1f, Klri2, Ncr1*, and *Tmsb15a*) from lungs (data available for download at *ArrayExpress* (*40, 49*) (accession E-MTAB-764) and available under a Creative Commons Attribution license. C57BL/6 mice were infected with influenza A/Puerto Rico/8/34.

### Data from Influenza Infection in *Ifnar1*^-/-^ Mice

We used published viral titers, IFN-γ, IFN-γ^+^ CD4^+^ and CD8^+^ T cells, and weight loss data digitized from a study where C57BL/6 or IFN-α receptor knock-out (*Ifnar1*^-/-^) mice were infected with influenza A/Fort Monmouth/1/47 (*36*).

### Data from Experimental Influenza Challenge in Humans

We used published viral titer, CD8^+^ T cell, and IFN-γ data from a study of adult human volunteers challenged with influenza A/California/04/2009 (data available for download from Rahil et al. (*42*)). Viral loads (copies/ml) were quantified from daily nasal washes. Activated (CD38^+^) effector (CD45RA^+^CD27^-^CCR7^-^) and memory (CD45RA^-^CD27^+^) CD8^+^ T cell frequencies and IFN-γ were quantified from collected blood. Only participants with sustained viral shedding were included. Missing or zero viral load values at early timepoints were imputed as previously described (*38*).

### Proteomics and Transcriptomics Data from Influenza Infection in Humans

We used published transcriptomic (*IFNG* and *IFNGR2*) and proteomic (IFNG) data from serum from a study of patients with moderate (non-ICU) or severe (ICU) influenza (*9, 10*). All data are available in public databases. Normalized transcript expression data are available from FigShare (10, 50) (https://doi.org/10.6084/m9.figshare.24299152), and proteome expression values are available from FigShare (9, 51) (https://doi.org/10.6084/m9.figshare.27826857).

### Mechanistic Mathematical Model of IFN-γ and IFN-γ^+^ Cell Regulation

We incorporated IFN-γ and IFN-γ^+^ cells into our mechanistic mathematical model describing CD8^+^ T cell-mediated clearance during influenza A virus infection (*6*). Briefly, our published model assumes that the virus (*V*) infects susceptible epithelial cells (*T*) at a rate β*V* day^-1^. Infected cells enter an eclipse phase (*I*_1_) before becoming virus-producing infected cells (*I*_2_) at a rate *k* day^-1^. Infected cells are cleared by innate mechanisms at rate δ day^-1^ and by CD8^+^ T cells in a density dependent manner 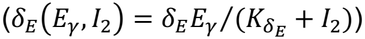. Virus is produced at a rate *p* TCID_50_ cell^-1^ day^-1^ and is cleared at a rate *c* day^-1^. The dynamics of effector CD8^+^ T cells, which are comprised predominantly of IFN-γ^+^ T cells (Fig. S1C), are described below, and a proportion of effector T cells take on a memory phenotype at rate ζ cell^-1^ day^-1^ after τ_*M*_ days. Total memory CD8^+^ T cells are defined by Ê_*M*_ = *E*_*M*_ + *E*_*M*0_, where *E*_*M*0_ is the initial number of memory CD8^+^ T cells.

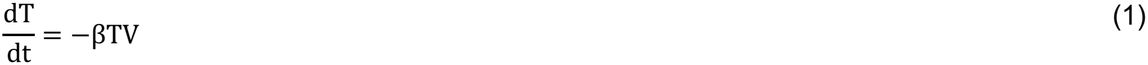

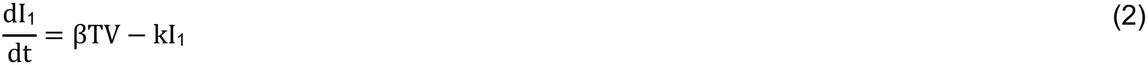

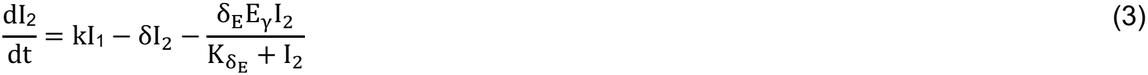

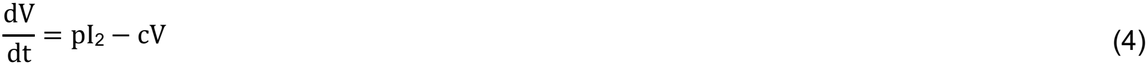

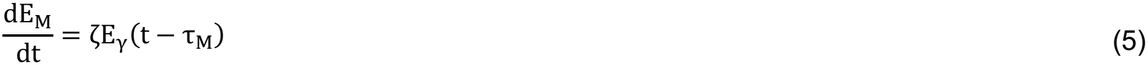

To describe the dynamics of IFN-γ, we developed equations detailing the activation and regulation of IFN-γ-producing cells. NK cells become activated and begin producing IFN-γ (*N*_γ_) proportional to eclipse cells at a rate ω_*N*_ cell^-1^ day^-1^ after τ_*N*_ days, which may reflect cross talk with dendritic cells or macrophages (*25, 52*). These cells are then downregulated by IFN-γ^+^ CD4^+^ T cells at a rate *g*_*N*_ (CD4^+^ T cells)^-1^ day^-1^. NK cell production of IFN-γ (*F*_*N*_) occurs at rate λ_*N*_ with downregulation by IFN-γ^+^ CD4^+^ T cells 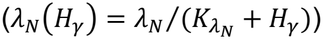. IFN-γ binds to NK cells at a rate µ cell^-1^ day^-1^ and to other cells at rate *q*_*N*_(*H*_γ_) = *q*_*N*_*H*_γ_/(*K*_q*N*_ + *H*_γ_) day^-1^. Natural decay occurs at a rate *d*_γ_ day^-1^.

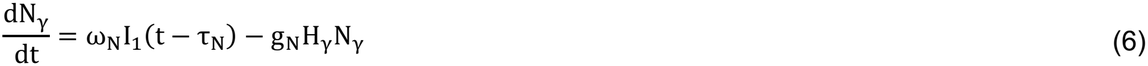

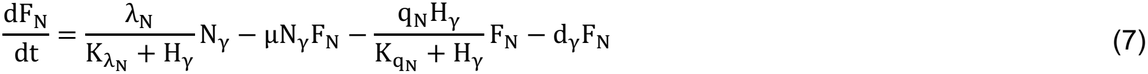

The activation of IFN-γ^+^ CD4^+^ T cells (*H*_γ_) and IFN-γ^+^ CD8^+^ T cells (*E*_γ_) is modeled using a linear chain trick with activation by IFN-γ^+^ NK cells (*20, 46, 47*) at rates ω_*H*_ and ω_*E*_ (NK cell)^-1^ day^-1^, respectively. Each T cell phenotype transitions through differentiation states at rates *a*_*H*_ = *n*_*H*_/τ_*H*_ and *a*_*E*_ = *n*_*E*_/τ_*E*_ day^-1^, where *n*_*H*_ and *n*_*E*_ are the number of states and τ_*H*_ and τ_*E*_ are the time delays, respectively. IFN-γ^+^ CD4^+^ and CD8^+^ T cells are downregulated by memory T cells at nonlinear rates 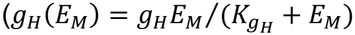 and 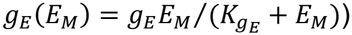, where *g*_*H*_ and *g*_*E*_ are the maximum rates, and 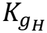 and 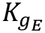 are the half-saturation constants. This may reflect a direct interaction of memory T cells or other immune responses occurring at the same time (*1, 24, 53*). We tested other mechanisms of downregulation, including linear self-regulation (*g*_*H*_(*E*_*M*_) = *g*_*H*_; *g*_*E*_(*E*_*M*_) = *g*_*E*_) or downregulation by memory CD8^+^ T cells (*g*_*H*_(*E*_*M*_) = *g*_*H*_*E*_*M*_; *g*_*E*_(*E*_*M*_) = *g*_*E*_*E*_*M*_), but these terms were unable to capture IFN-γ^+^ T cells kinetics or their fluorescence activity. We also tested the addition of a term describing IFN-γ^+^ CD4^+^ T cell activation of IFN-γ^+^ CD8^+^ T cells, but this added complexity was not supported by the data.

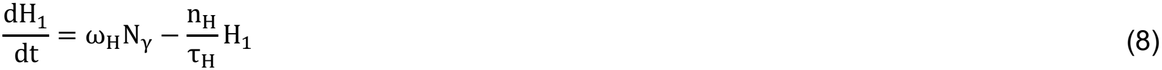

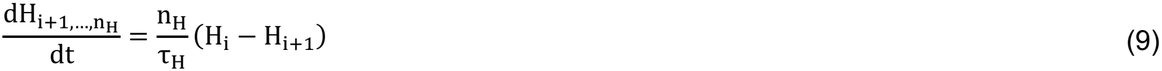

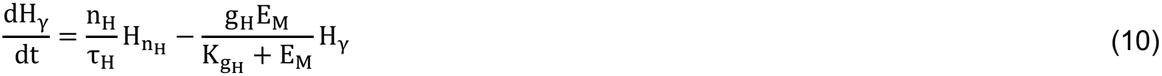

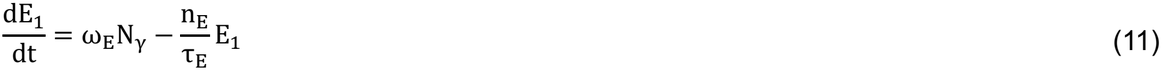

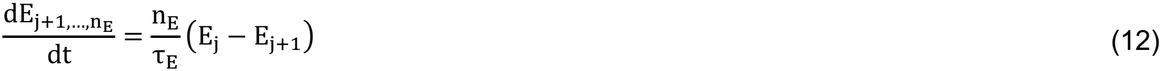

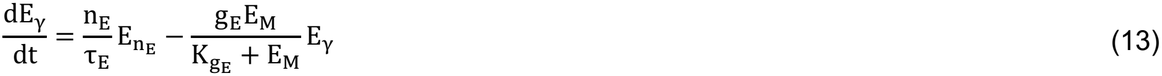

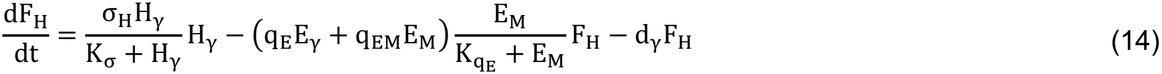

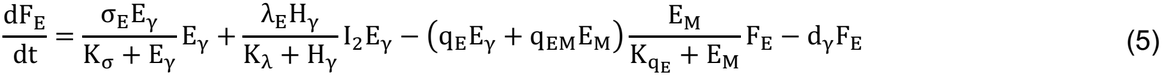

Because multiple factors have been shown to regulate IFN-γ from T cells, including various cytokines and interactions with antigen-presenting cells (*1, 14, 16, 53–55*), the following regulatory terms were included in the model. Regulation of IFN-γ production by CD4^+^ T cells occurs as a functional response according to λ_*E*_(*H*_γ_) = λ_*E*_*H*_γ_/(*K*_λ_ + *H*_γ_). Self-regulation of IFN-γ production was required for both CD4^+^ and CD8^+^ T cells and described by the nonlinear rates σ_*H*_(*H*_γ_) = σ_*H*_ *H*_γ_/(*K* _*σ*_ + *H*_γ_) and σ_*E*_(*E*_γ_) = σ_*E*_ *E*_γ_/(*K*_*σ*_ + *E*_γ_), which may occur via concurrent IL-10 production(*1, 53, 56*). Decay of T cell-derived IFN-γ occurs at a rate *d*_γ_ day^-1^, and uptake of IFN-γ by all responding CD8^+^ T cells (*E*_γ_ + *E*_*M*_) occurs at the nonlinear rates 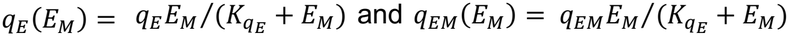 and *q*_*EM*_(*E*_*M*_) = *q*_*EM*_*E*_*M*_/(*K*_*qE*_ + *E*_*M*_). Total IFN-γ is defined by *F* = *F*_*N*_ + *F*_*H*_ + *F*_*E*_ + *F*_0_, where *F*_0_ is the initial concentration of IFN-γ.

It was necessary to model the fluorescence intensity to constrain IFN-γ production from each cell type. To model this, the IFN-γ production terms for IFN-γ^+^ NK cells, IFN-γ^+^ CD4^+^ T cells, and IFN-γ^+^ CD8^+^ T cells were multiplied by a scaling factor (ε) to translate the scale (iMFI_N_ = ελ_N_(H_*γ*_)N_*γ*_; iMFI_H_ = ε(σ_H_(H_*γ*_)H_*γ*_); and iMFI_E_ = ε(σ_E_(E_*γ*_)E_*γ*_ + λ_E_(H_*γ*_ (I_2_E_*γ*_), respectively).

### Parameter Quantification for Murine Infections

To quantify the model parameters, the model was simultaneously fit to viral loads, IFN-γ, IFN-γ^+^ NK cells, IFN-γ^+^ CD4^+^ T cells, IFN-γ^+^ CD8^+^ T cells, memory CD8^+^ T cells, and the integrated median fluorescent intensity using a simulated annealing global optimization algorithm as previously described (*6, 57*) using MATLAB 2025b (*58*) and delayed differential equation solvers *dde23* or *dde15s*. To account for the lower limit of detection (LOD) of IFN-γ^+^ cells, model outputs below the LOD were set to the value at day 0. All model parameters were estimated except for the number of states in the linear chains (*n*_*E*_ and *n*_*H*_), which were fixed to 20. The initial number of target cells (*T*(0)) was set to 1 × 10^7^ cells(*6, 44, 57*). The initial number of eclipse cells was set to *I*_1_(0) = 75 (*6, 44, 57*). The initial virus (*V*(0)), memory CD8^+^ T cells (*E*_*M*_(0))) (*6, 44, 57*), IFN-γ^+^ NK cells (*N*_γ_(0)), IFN-γ^+^ CD4^+^ T cells (*H*_γ_(0)), IFN-γ^+^ CD8^+^ T cells (*E*_γ_(0)) and IFN-γ (*F*(0)) were set to zero.

Because our prior work showed that parameter adjustments were needed to account for changes induced by the CD8 depletion antibody (*6*), the best-fit values from fitting the control infection were systematically varied until the viral load, IFN-γ, and imputed levels of IFN-γ^+^ CD4^+^ T cells, IFN-γ^+^ CD8^+^ T cells, and memory CD8^+^ T cells could be simultaneously fit. Because IFN-γ^+^ T cell data was not available from the depletion experiment and the concordance between IFN-γ^+^ T cells and total T cells from the control experiment, IFN-γ^+^ CD4^+^, IFN-γ^+^ CD8^+^, and memory CD8^+^ T cells were assumed to change proportionally to the total.

The best-fit values from the IFN-γ model were also used to simulate infection in C57BL/6 control and *Ifnar1*^-/-^ murine with parameter adjustments required to account for differences in murine model as described in the text. Instead of fitting to data, model outputs were matched to data according to a scaling factor to account for differences in data acquisition and data quantification (0.93 for viral loads, 0.69 for IFN-γ, and 0.84 for IFN-γ^+^ CD4^+^ and CD8^+^ T cells).

### Parameter Quantification for Human Influenza Challenge

To model IFN-γ responses during human challenge infections, the parameters from our previous study were adapted as follows (*38*). Select parameters were held fixed (*E*_0_, *EM*_0_, *T*_0_, β, *p, c, k*, δ, δ_*E*_, *K*_*D*_, ζ, τ_*M*_, and *n*_*E*_), and the parameters describing IFN-γ^+^ CD8^+^ T cell responses (ω_*N*_, *g*_*N*_, τ_*N*_, ω_*E*_, τ_*E*_, *g*_*E*_, and *K*_*g*E_) were estimated. Because the data was insufficient to estimate parameters related to IFN-γ^+^ CD4^+^ T cells, it was assumed that these cells behave similarly to IFN-γ^+^ CD8^+^ T cells and were scaled according to the parameter values obtained during the murine fits: *n*_*H*_ = *n*_*E*_, ω_*H,i*_ = ω_*H,M*_ ω_*E,i*_ /ω_*E,M*_, τ_*H,i*_ = τ_*H,M*_ τ_*E,i*_ /τ_*E,M*_, *g*_*H,i*_ = *g*_*H,M*_ *g*_*E,i*_ /*g*_*E,M*_, and *K*_*gH,i*_ = *K*_*gH,M*_ *K*_*gE,i*_ /*K*_*gE,M*_, where subscript *i* denotes patient *i* and *M* denotes the murine value. This approach to parameterization allowed for the IFN-γ^+^ NK cell kinetics to be predicted.

To quantify serum IFN-γ, the IFN-γ measurements from two patients (ID no. 111 and 307) that exhibited sufficient changes from baseline were used. Because these measurements came from serum rather than lung, and the data could not constrain all the production and uptake parameters, we identified a minimal set that reproduced the timing and relative magnitude of the IFN-γ response. Because there were increased numbers of human CD8^+^ T cells compared to murine CD8^+^ T cells, the T cell half-saturation constants were scaled by the best-fit IFN-γ^+^ CD8^+^ T cell regulation parameter, *K*_*gE*_; *K*_*qN,i*_ = 10^1.5^*K*_*qN,M*_ *K*_*gE,i*_/*K*_*gE,M*_, *K*_λ,*i*_ = 10^1.5^*K*_λ,*M*_ *K*_*gE,i*_/*K*_*gE,M*_, *K*_σ,*i*_ = 10^1.5^*K*_σ,*M*_ *K*_*gE,i*_/*K*_*gE,M*_, and *K*_*qE,i*_ = 10^1.5^*K*_*qE,M*_ *K*_*gE,i*_/*K*_*gE,M*_. IFN-γ^+^ T cell production and receptor-mediated uptake were scaled by the best-fit differences in IFN-γ^+^ T cell-mediated infected cell clearance rate (δ_*E*_) and memory formation rate (ζ); *q*_*E,i*_ = *q*_*E,M*_ δ_*E,i*_/δ_*E,M*_, *q*_*EM,i*_ = *q*_*EM,M*_ δ_*E,i*_/δ_*E,M*_, σ_*H,i*_ = 10^0.5^σ_*H,M*_ (δ_*E,i*_ + ζ_,*i*_)g(δ_*E,M*_ + ζ_,*M*_), and σ_*E,i*_ = 10^0.5^σ_*E,M*_ (δ_*E,i*_ + ζ_,*i*_)g(δ_*E,M*_ + ζ_,*M*_). Finally, to ensure NK cell-derived IFN-γ peaks were below T cell-derived IFN-γ peaks, the uptake was scaled by the best-fit change in IFN-γ^+^ T cell activation (ω_*E*_): µ_,*i*_ = 10^0.75^µ_,*M*_ ω_*E,i*_/ω_*E,M*_, and *q*_*N,i*_ = 10^0.75^*q*_*N,M*_ ω_*E,i*_/ω_*E,M*_. The constant factors (10^1.5^, 10^0.5^, 10^0.75^) were selected to reproduce the qualitative timing and relative magnitude of the human IFN-γ response. Separate axes were used to display serum measurements and modeled lung kinetics because their magnitudes were not directly comparable. These rules were then applied to the other patients in the cohort to infer their IFN-γ kinetics.

### Virtual Patient Cohorts for Mild, Moderate, and Severe Influenza

Virtual patient simulations for mild, moderate, and severe influenza were performed by sampling parameters from normal distributions with mean and standard deviations taken from parameter sets used for the human challenge data. Parameter combinations were rejected according to the following criteria.

Rejected viral load trajectories included those that (i) were too low as defined by those that failed to surpass 90% of the minimum human challenge viral titer data during days 1-3 pi, (ii) were too high as defined by those that exceeded 1 × 10^9^ TCID_50_, or (iii) did not clear within 10 dpi. IFN-γ^+^ CD4^+^ and CD8^+^ T cell trajectories were required to stay below 5 × 10^5^ cells at 1 dpi and 1 × 10^7^ at 2 dpi to limit premature expansion and were rejected if they were below 90% of the minimum human challenge data from 1-6 dpi or if they remained above 10% of the maximum digital twin prediction at 10 dpi to ensure T cell clearance post-infection. Memory CD8^+^ T cell trajectories were required to have values above 90% of the minimum human challenge data during days 1-6 pi. Additional criteria included consistency with qualitative features of the human challenge and murine datasets, where CD8^+^ T cell peaks were required to occur around the time of viral clearance (2 d before or 1 d after), CD4^+^ T cell peaks were required to occur within 1 d of the CD8^+^ T cell peak and within 1 log_10_ of the CD8^+^ T cell magnitude, IFN-γ peaks were required to occur before the CD8^+^ T cell peak., and IFN-γ^+^ NK cells were required to turn off production by 6 dpi and peak between 1.4 and 2.7 days (Fig. S6).

Virtual patients were stratified into model-based severity groups according to the cumulative area under the curve (CAUC) of infected cells, which we previously established to correlate with acute lung injury (*6*). Virtual patients with their CAUC less than the maximum CAUC obtained from the model fits to the human challenge data were classified as mild influenza. Virtual patients with CAUC > 1 × 10^9^ were classified as severe influenza (ICU), and the remaining virtual patients were classified as having moderate influenza (non-ICU). Because the virtual patients lacked observed clinical outcomes, these groups represent model-derived severity strata rather than clinical classifications.

### Statistical Analyses

Unpaired *t* tests were used to assess significance. *P* < 0.05 was considered significant. Alternate model formulations were compared using the Akaike Information Criteria (AIC).

## Author Contributions

Conceptualization: A.M.S.

Methodology: A.M.S., A.P.S., C.J.S., N.B., H.S.H., M.H.,

Investigation: A.M.S., A.P.S., C.J.S.

Visualization: A.M.S., C.J.S. Supervision: A.M.S.

Writing—original draft: A.M.S., A.P.S., C.J.S.

Writing—review & editing: A.M.S., C.J.S., N.B., K.S., M.H., H.S.H., A.P.S.

## Data and Code Availability

Newly generated data needed will be available upon publication. Original published murine transcriptomics data are available from *ArrayExpress* (*40, 49*) (accession E-MTAB-764); normalized human transcript expression data are available from FigShare (10, 50) (https://doi.org/10.6084/m9.figshare.24299152); normalized, human proteome expression data are available from FigShare (9, 51) (https://doi.org/10.6084/m9.figshare.27826857).

