## Supplemental Figures and Tables for "The IFN-γ Paradox: Mechanistic Modeling Uncouples Cytokine Abundance from Influenza Pathogenesis"

### Supplementary Figures

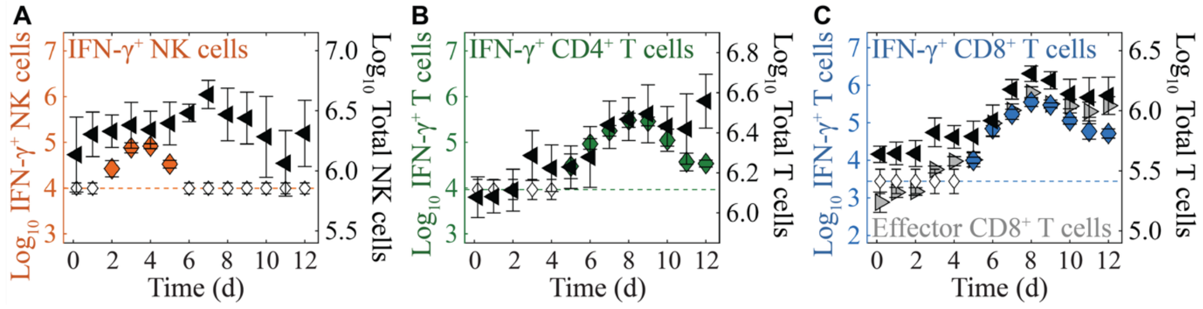


**Fig. S1. Proportion of IFN-γ$^+$ NK cells, CD4$^+$ T cells, and CD8$^+$ T cells during Influenza A Virus Infection in BALB/c mice.** (**A**) IFN-γ$^+$ NK cells (orange, left axis) are a small proportion of the broader NK cell population (black, right axis) and therefore cannot be inferred from the total number. (**B**-**C**) IFN-γ$^+$ CD4$^+$ T cells (green, left axis) and IFN-γ$^+$ CD8$^+$ T cells (blue, left axis) make up a significant proportion of their broader populations (black, right axes). IFN-γ$^+$ CD8$^+$ T cells also have similar dynamics as effector CD8$^+$ T cells (CD25$^-$CD43$^+$) (*6*), supporting their functionality.

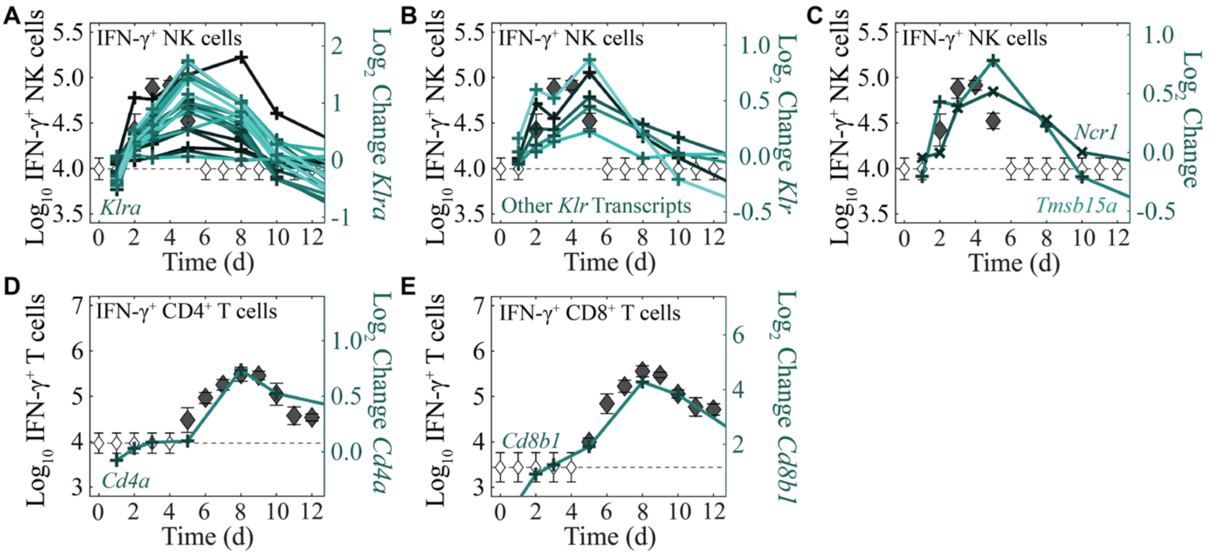


**Fig. S2. Comparison of IFN-γ$^+$ NK cells and IFN-γ$^+$ T cells with Transcriptomic Data.** (**A-C**) Transcripts previously linked to NK cells (*38*), including expression of (**A**) *Klra1-Klra23*; (**B**) *Klrb1a*, *Klrb1c*, *Klre1*, *Klrb1f*, and *Klri2*; and (**C**) *Ncr1* and *Tmsb15a* did not align with IFN-γ$^+$ subsets of NK cells. (**D**) *Cd4a* expression was similar to IFN-γ$^+$ CD4$^+$ T cells. (**E**) Similar to *Cd8a (Fig. 1D)*, *Cd8b1* expression reflected IFN-γ$^+$ CD8$^+$ T cells.

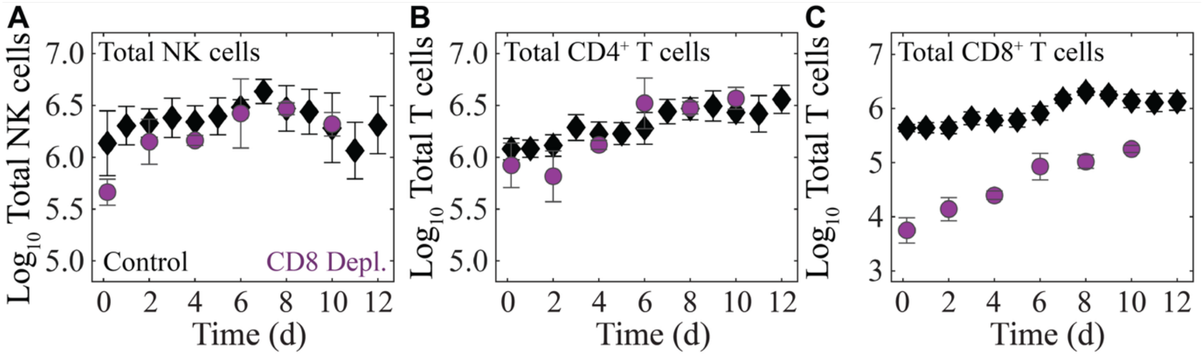


**Fig. S3. NK cells, $CD4^+$ T cells, and $CD8^+$ T cells following $CD8^+$ T cell Depletion.** (**A**-**C**) The dynamics of total NK cells (**A**), $CD4^+$ T cells (**B**), and $CD8^+$ T cells (**C**) in control (black) and $CD8^+$ T cell-depleted mice infected with influenza A/PR8 (*6*). NK cells and $CD4^+$ T cells were significantly different during early infection (0 and 2 dpi, respectively) but then recovered to the level of the control animals. $CD8^+$ T cells were significantly lower throughout.

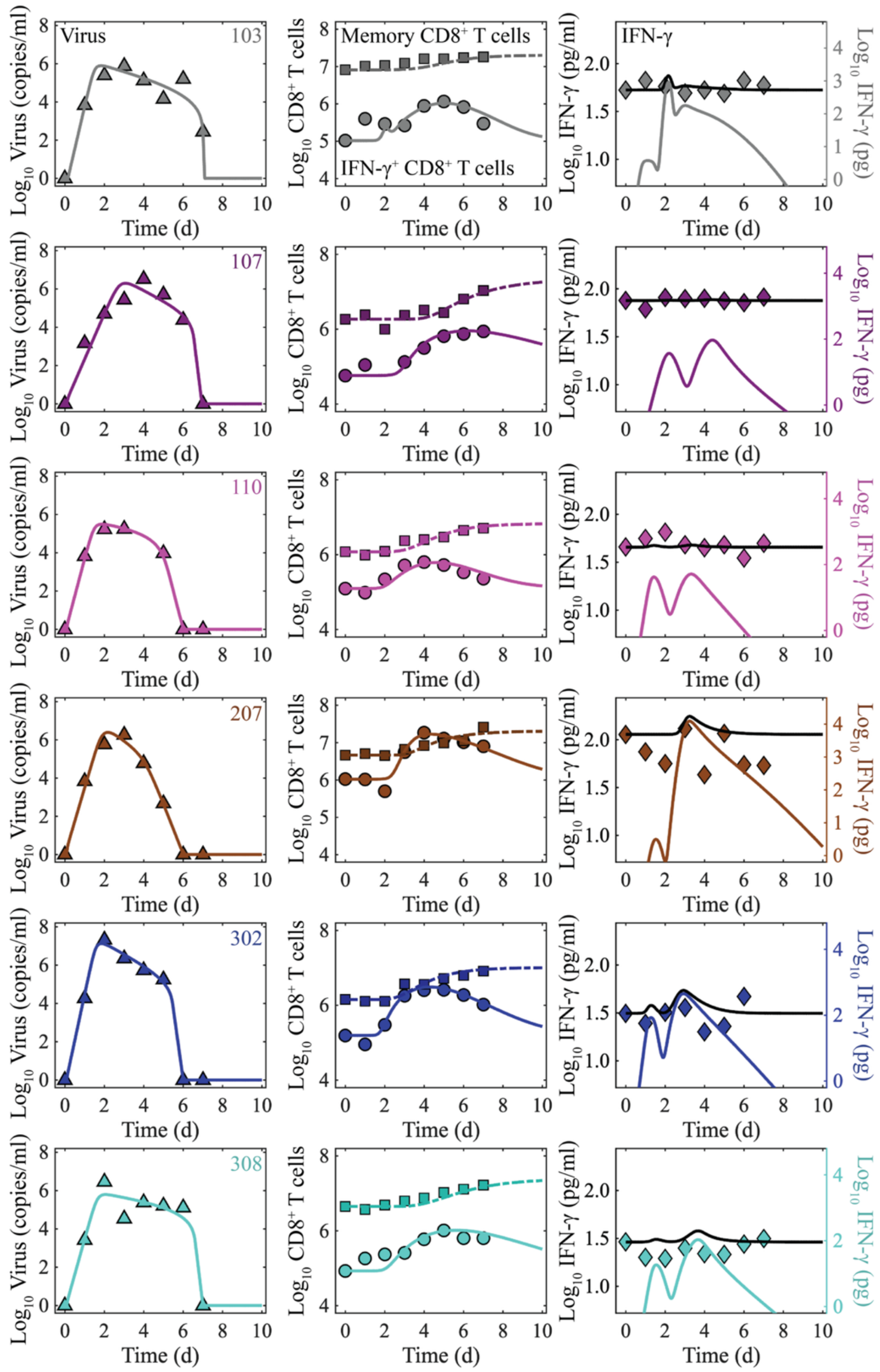


**Fig. S4. Model Fits to Human Influenza Challenge Data.** Several patients experimentally challenged with influenza A/California/04/2009 virus did not have significant increases in IFN-γ in the serum (colored diamonds, right axes) (*40*). Model simulations (black, left axes) for viral loads (triangle), effector CD8$^+$ T cells (circle), and memory CD8$^+$ T cells (square) for each individual patient. Parameter translations across the cohort are described in the main text and Table S3.

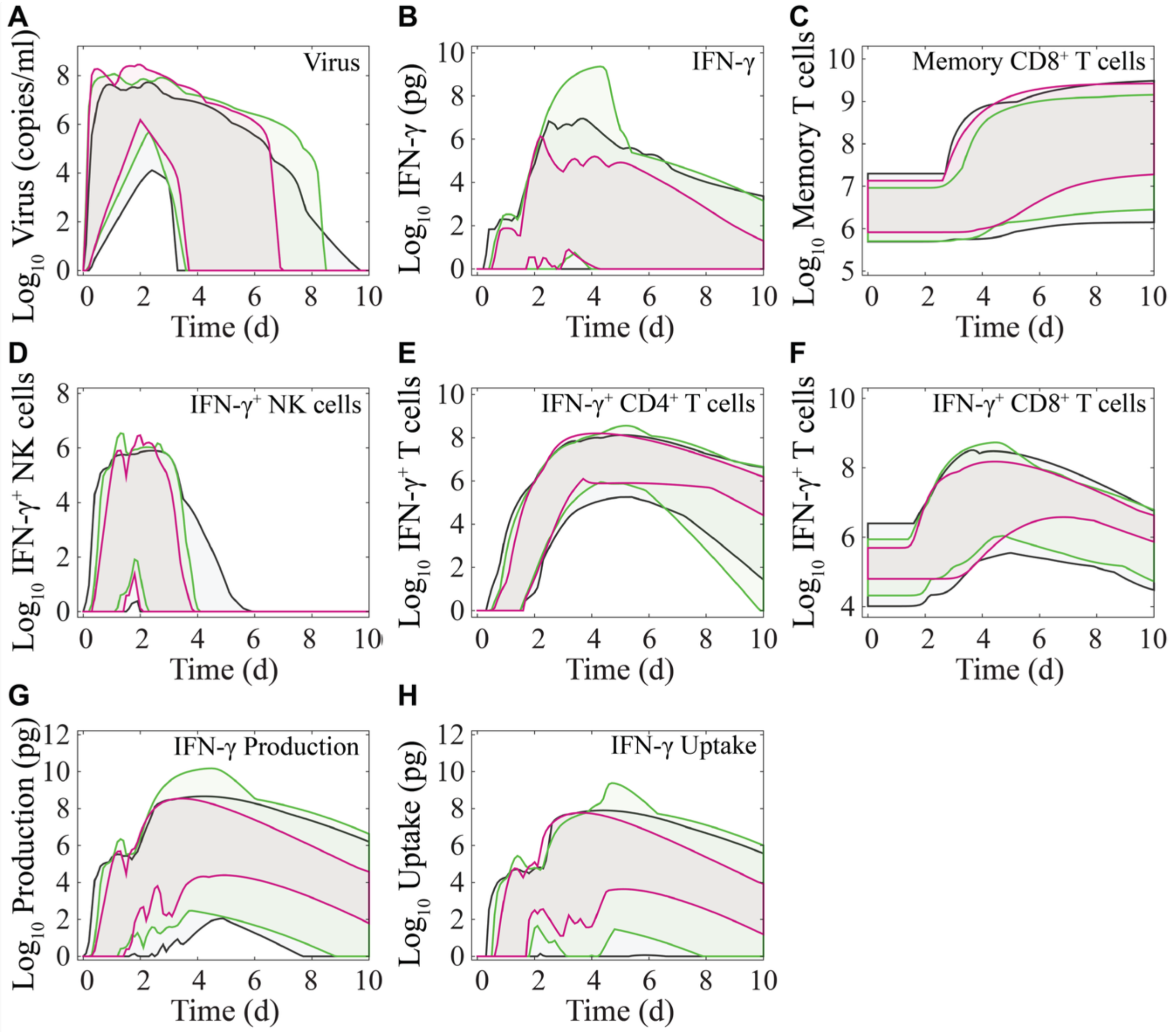


**Fig. S5: Virtual Patient Cohorts for Mild, Moderate, and Severe Influenza**. Virtual patients were generated and stratified by severity level (mild [grey], non-ICU [green], and ICU [pink] cohorts) as described in the main text. The upper and lower bounds of each cohort's kinetics for each cohort showed significant overlap. The minimum and maximum values of (**A**) virus, (**B**) IFN-γ, (**C**) memory CD8$^+$ T cells, (**D**) IFN-γ$^+$ NK cells, (**E**) IFN-γ$^+$ CD4$^+$ T cells, (**F**) IFN-γ$^+$ CD8$^+$ T cells, (**G**) IFN-γ production, and (**H**) receptor-mediated IFN-γ uptake kinetics are shown.

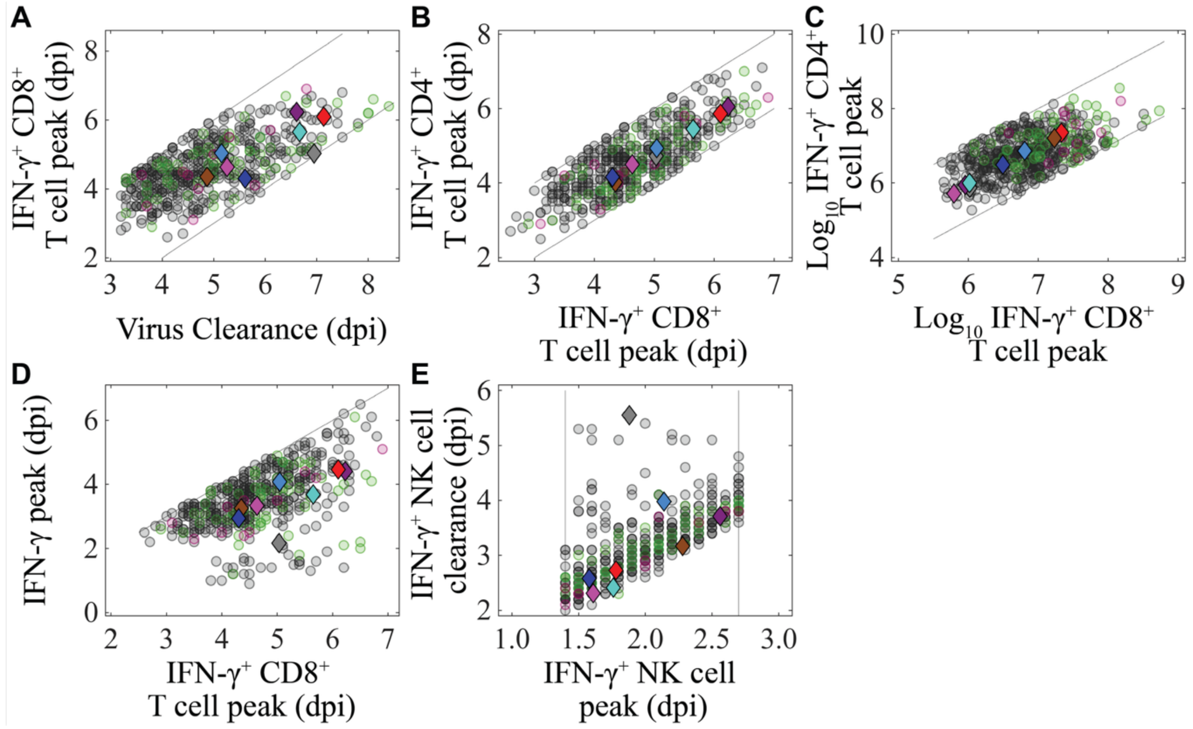


**Fig. S6: Feature Selection Used to Generate Realistic Virtual Patient Trajectories.** Kinetic features of the virtual patients (circles) across mild (grey), non-ICU (green), and ICU (magenta) cohorts compared with those from the personalized trajectories for human patients challenged with influenza (diamonds, colors matched to patients in Fig. 3 and Fig. S4). Virtual patients were retained if their (**A**) IFN-γ$^{+}$ CD8$^{+}$ T cell peaks occurred between one day before and two days after the virus clearance; (**B-C**) IFN-γ$^{+}$ CD4$^{+}$ T cell peaks occurred within one day of the IFN-γ$^{+}$ CD8$^{+}$ T cell peak and within 1 $\log_{10}$ of the IFN-γ$^{+}$ CD8$^{+}$ T cell peak; (**D**) IFN-γ peaked before the IFN-γ$^{+}$ CD8$^{+}$ T cell peak; and (**E**) IFN-γ$^{+}$ NK cells peaked between 1.4 and 2.7 dpi and cleared before 6 dpi.

**Supplementary Tables**

**Table S1: IFN-γ Model Parameters.** Model variable, description, best fit value, and 95% confidence interval associated with model fits in Figure 1.

| Parameter | Description | Units | Value | CI |
|---|---|---|---|---|
| $p$ | Virus production | $TCID_{50}$ cell$^{-1}$ d$^{-1}$ | 1.2 | [7.0 × $10^{-1}$, 5.2] |
| $c$ | Virus clearance | d$^{-1}$ | 11.0 | [6.1, 29.0] |
| $\beta$ | Virus infectivity | $TCID_{50}^{-1}$ d$^{-1}$ | 5.5 × $10^{-5}$ | [1.4 × $10^{-5}$, 8.7 × $10^{-5}$] |
| $k$ | Eclipse phase transition | d$^{-1}$ | 4.0 | [4.0, 5.9] |
| $\delta$ | Infected cell clearance | d$^{-1}$ | 3.0 × $10^{-1}$ | [2.1 × $10^{-1}$, 4.9 × $10^{-1}$] |
| $\delta_E$ | Infected cell clearance by effector CD8$^+$ T cells | Cells $E_\gamma^{-1}$ d$^{-1}$ | 9.4 | [1.8, 11.0] |
| $K_d$ | Half-saturation constant | Cells | 2.5 × $10^3$ | [1.0 × $10^2$, 3.4 × $10^5$] |
| $\zeta$ | Memory CD8$^+$ T cell generation | $E_M$ $E_\gamma^{-1}$ d$^{-1}$ | 2.2 × $10^{-2}$ | [1.1 × $10^{-2}$, 8.7 × $10^{-2}$] |
| $\tau_M$ | Memory CD8$^+$ T cell generation delay | d | 2.0 | [2.0, 3.8] |
| $\omega_N$ | IFN-γ$^+$ NK cell activation | $N_\gamma$ cells$^{-1}$ d$^{-1}$ | 2.9 × $10^{-2}$ | [2.3 × $10^{-2}$, 6.7 × $10^{-2}$] |
| $g_N$ | IFN-γ$^+$ NK cell downregulation | $H_\gamma^{-1}$ d$^{-1}$ | 8.0 × $10^{-5}$ | [3.0 × $10^{-5}$, 2.7 × $10^{-4}$] |
| $\tau_N$ | IFN-γ$^+$ NK cell delay | d | 5.8 × $10^{-1}$ | [2.1 × $10^{-1}$, 8.0 × $10^{-1}$] |
| $\omega_H$ | IFN-γ$^+$ CD4$^+$ T cell activation | $H_\gamma$ $N_\gamma^{-1}$ d$^{-1}$ | 1.5 | [5.4 × $10^{-1}$, 2.9] |
| $n_H$ | IFN-γ$^+$ CD4$^+$ T cell chain links | 1 | 20 | - |
| $\tau_H$ | IFN-γ$^+$ CD4$^+$ T cell activation delay | d | 3.2 | [2.6, 3.7] |
| $g_H$ | IFN-γ$^+$ CD4$^+$ T cell downregulation | d$^{-1}$ | 1.4 | [3.5 × $10^{-1}$, 2.8] |
| $K_{g_H}$ | Half-saturation constant | $E_M$ | 8.5 × $10^3$ | [1.1 × $10^2$, 4.6 × $10^4$] |
| $\omega_E$ | IFN-γ$^+$ CD8$^+$ T cell activation | $E_\gamma$ $N_\gamma^{-1}$ d$^{-1}$ | 1.5 | [5.0 × $10^{-1}$, 2.4] |
| $n_E$ | IFN-γ$^+$ CD8$^+$ T cell chain links | 1 | 20 | - |
| $\tau_E$ | IFN-γ$^+$ CD8$^+$ T cell activation delay | d | 3.5 | [2.9, 3.9] |
| $g_E$ | IFN-γ$^+$ CD8$^+$ T cell downregulation | d$^{-1}$ | 9.6 × $10^{-1}$ | [2.9 × $10^{-1}$, 3.0] |
| $K_{g_E}$ | Half-saturation constant | $E_M$ | 6.1 × $10^3$ | [1.1 × $10^2$, 4.7 × $10^4$] |
| $\lambda_N$ | NK cell IFN-γ production | pg $H_\gamma$ $N_\gamma^{-1}$ d$^{-1}$ | 3.0 × $10^3$ | [1.2 × $10^3$, 5.0 × $10^4$] |
| $K_{\lambda_N}$ | Half-saturation constant | $H_\gamma$ | 4.6 × $10^3$ | [9.1 × $10^2$, 9.1 × $10^4$] |
| $d_\gamma$ | IFN-γ clearance | pg d$^{-1}$ | 5.7 | [4.0, 46.0] |
| $K_\lambda$ | Half-saturation constant | $H_\gamma$ | 7.0 × $10^4$ | [6.8 × $10^3$, 5.0 × $10^5$] |
| $\sigma_H$ | CD4$^+$ T cell IFN-γ production | $H_\gamma^{-1}$ d$^{-1}$ | 2.4 × $10^{-1}$ | [1.5 × $10^{-1}$, 1.3] |
| $K_\sigma$ | Half-saturation constant | T cells | 6.3 × $10^3$ | [1.8 × $10^3$, 3.4 × $10^5$] |
| $\lambda_E$ | CD8$^+$ T cell IFN-γ production promoted by CD4$^+$ T cells | $E_\gamma^{-1}$ cells$^{-1}$ d$^{-1}$ | 1.6 × $10^{-7}$ | [6.0 × $10^{-8}$, 8.0 × $10^{-6}$] |
| $\sigma_E$ | CD8$^+$ T cell IFN-γ production | $E_\gamma^{-1}$ d$^{-1}$ | 5.3 × $10^{-1}$ | [2.5 × $10^{-1}$, 2.4] |
| $\mu$ | NK cell IFN-γ uptake | $N_\gamma^{-1}$ d$^{-1}$ | 1.9 × $10^{-4}$ | [4.4 × $10^{-5}$, 9.7 × $10^{-4}$] |
| $q_N$ | CD4$^+$ T cell IFN-γ uptake | d$^{-1}$ | 24.0 | [6.0 × $10^{-1}$, 6.5 × $10^2$] |
| $K_{q_N}$ | Half-saturation constant | $N_\gamma$ | 1.7 × $10^3$ | [1.0 × $10^2$, 4.8 × $10^5$] |
| $q_E$ | Effector CD8$^+$ T cell IFN-γ uptake | $E_\gamma^{-1}$ d$^{-1}$ | 3.1 × $10^{-4}$ | [6.7 × $10^{-5}$, 7.0 × $10^{-3}$] |
| $q_{E_M}$ | Memory CD8$^+$ T cell IFN-γ uptake | $E_M^{-1}$ d$^{-1}$ | 9.4 × $10^{-3}$ | [4.3 × $10^{-3}$, 7.9 × $10^{-2}$] |
| $K_{qE}$ | Half-saturation constant | $E_M$ | 3.1 × $10^2$ | [1.2 × $10^2$, 3.9 × $10^4$] |
| $\epsilon$ | Production to iMdFI scaling | 1 | 20.0 | [6.3 × $10^{-3}$, 4.0 × $10^{-2}$] |
| $E_{M0}$ | Memory CD8$^+$ T cell baseline | $E_M$ | 1.0 × $10^7$ | [7.0 × $10^2$, 9.0 × $10^2$] |
| $F_0$ | IFN-γ baseline | pg | 2.6 × $10^{-2}$ | [11, 35] |
| $T(0)$ | Initial uninfected cells | Cells | 1.0 × $10^7$ | - |
| $I_1(0)$ | Initial eclipse cells | Cells | 75 | - |

**Table S2: Model Parameters for CD8$^+$ T cell Depletion.** Comparison of parameters from fitting the model to control and CD8$^+$ T cell-depleted data. All other model parameters were unchanged from the control (Table S1).

| Parameter | Description | Value | |
|---|---|---|---|
| | | Control | CD8 Depl. |
| $T(0)$ | Initial uninfected cells | $1.0 \times 10^{7}$ | $4.2 \times 10^{6}$ |
| $\tau_N$ | IFN-γ$^+$ NK cell delay | $5.8 \times 10^{-1}$ | $1.5 \times 10^{-1}$ |
| $\omega_H$ | IFN-γ$^+$ CD4$^+$ T cell activation | 1.5 | 5.6 |
| $\tau_H$ | IFN-γ$^+$ CD4$^+$ T cell activation delay | 3.2 | 2.9 |
| $K_{g_H}$ | Half-saturation constant | $8.5 \times 10^{3}$ | $7.0 \times 10^{2}$ |
| $\tau_E$ | IFN-γ$^+$ CD8$^+$ T cell activation delay | 3.5 | 4.3 |
| $K_{g_E}$ | Half-saturation constant | $6.1 \times 10^{3}$ | $1.7 \times 10^{2}$ |

**Table S3: Model Parameters for *Ifnar1*$^{-/-}$ groups.** Comparison of parameters between BALB/c, C57BL/6 control, and *Ifnar1*$^{-/-}$ groups. All other model parameters were unchanged from the BALB/c control (Table S1).

| Parameter | Description | Value | | |
|---|---|---|---|---|
| | | BALB/c CTRL | C57BL/6 CTRL | C57BL/6 *Ifnar1*$^{-/-}$ |
| $\mu$ | NK cell IFN-γ uptake | $1.9 \times 10^{-4}$ | $2.0 \times 10^{-5}$ | $2.0 \times 10^{-5}$ |
| $T(0)$ | Initial uninfected cells | $1.0 \times 10^{7}$ | $7.0 \times 10^{6}$ | $2.0 \times 10^{7}$ |
